# Rainbow trout in the inlet tributaries of Lake Chinishibetsu, Shiretoko Peninsula

**DOI:** 10.1101/2023.03.23.533890

**Authors:** Genki Sahashi, Mari Kuroki, Takahiro Nobetsu, Kentaro Morita

## Abstract

Rainbow trout, Oncorhynchus *mykiss*, is one of the most widely introduced fish species in the world, and its impacts on native fishes and ecosystems are of considerable concern. One of the rivers inhabited by rainbow trout in the Shiretoko Peninsula is the Chinishibetsu River, and the origin of rainbow trout in this river is thought to be Lake Chinishibetsu in the upper reaches of the system, where the private stocking of rainbow trout was conducted in the 1960s. However, the basic biology of rainbow trout in the Lake Chinishibetsu area is currently unknown. This study addresses this knowledge gap by examining the biology of rainbow trout in the inlet tributaries of Lake Chinishibetsu based on sampling conducted during the rainbow trout spawning season. A total of 104 rainbow trout, ranging in age from 1+ to 8+ years, were collected from the two inlet tributaries of Lake Chinishibetsu. White-spotted charr *Salvelinus leucomaenis* and Siberian stone loach *Barbatula oreas*, neither of which is native to the Shiretoko Peninsula, were also collected and had presumably invaded the area at the same time as the rainbow trout. The sampled rainbow trout included immature and mature males and females. The distribution of fork lengths of mature females was bimodal, and the sex ratio of mature rainbow trout was male-biased. Our results indicate that the rainbow trout population in the inlet tributaries of Lake Chinishibetsu is reproducing continuously and exhibits a dimorphic life history with river residents and lake migrants of both sexes. Additionally, rainbow trout continue to be collected downstream of the Chinishibetsu River, which is the primary habitat for this species in the Shiretoko Peninsula. Therefore, unless rainbow trout are eliminated from Lake Chinishibetsu, which serves as a source of non-native species upstream of the Chinishibetsu River, it will be difficult to control rainbow trout distributions and minimize population sizes on the Shiretoko Peninsula.

## Introduction

Rainbow trout, *Oncorhynchus mykiss*, is one of the most widely introduced fish species in the world (Crawford & Muir, 2008), and its negative impacts on native fishes and ecosystems are of considerable concern (Dunham et al., 2004; Fausch, 2007; Stanković et al., 2015). The presence of introduced rainbow trout has been widely confirmed in Japan (Morita, 2019; Hasegawa, 2020), and the species has been reported to inhabit several rivers on the Shiretoko Peninsula (Morita et al., 2003; Yamamoto, 2008; Kasai et al., 2010). One of the rivers on the Shiretoko Peninsula where rainbow trout is distributed is the Chinishibetsu River, where rainbow trout is collected every year in the lower river reaches during monitoring surveys carried out by the Shiretoko World Natural Heritage Site (Hokkaido Regional Forest Office & Forest Realize Co., Ltd, 2022). The origin of the rainbow trout in the Chinishibetsu River is thought to be Lake Chinishibetsu in the upper reaches of the system, where private stocking of rainbow trout was conducted in the 1960s (Kaizaki, 1990). However, no basic biological information has been reported for the non-native trout population of Lake Chinishibetsu. This study examined the population density, distribution of body sizes, age composition, and maturity of rainbow trout in the inlet tributaries of Lake Chinishibetsu through fish sampling during the rainbow trout spawning season.

## Materials and methods

This study was conducted in May 2017 and April 2020 on two inlet tributaries of Lake Chinishibetsu (Fig. 1). The river flowing out of Lake Chinishibetsu has a natural waterfall near its confluence with the main stream that does not allow fish to migrate upstream (Fig. 1). Thus, any non-native fish found in the inlet tributaries of Lake Chinishibetsu are assumed to have been introduced artificially rather than naturally invading from other water sources. Fish sampling was conducted in May 2017 in Tributary A (river width 2.9 ± 1.1 m), and in April 2020 in Tributaries A and B (Figs. 1, 2). Fish were collected using a backpack electrofishing unit (200–300 V DC, model 12B or LR-20B; Smith-Root, Vancouver, WA, USA) and a dip net (30-cm width, a mesh size 3 mm). All salmonid fishes obtained during sampling were promptly frozen in the laboratory. After defrosting, the fork length (FL), sex, reproductive state (mature or immature), and age were determined for each individual (age was determined by using otoliths). For the fish sampling in 2017, a 40-m long study reach was established in Tributary A (Fig. 1), and rainbow trout densities in the reach were estimated using the two-pass removal method (model M[b], program CAPTURE, Rexstad & Burnham, 1991; https://www.mbr-pwrc.usgs.gov/software/capture.html).

**Figure 1.**
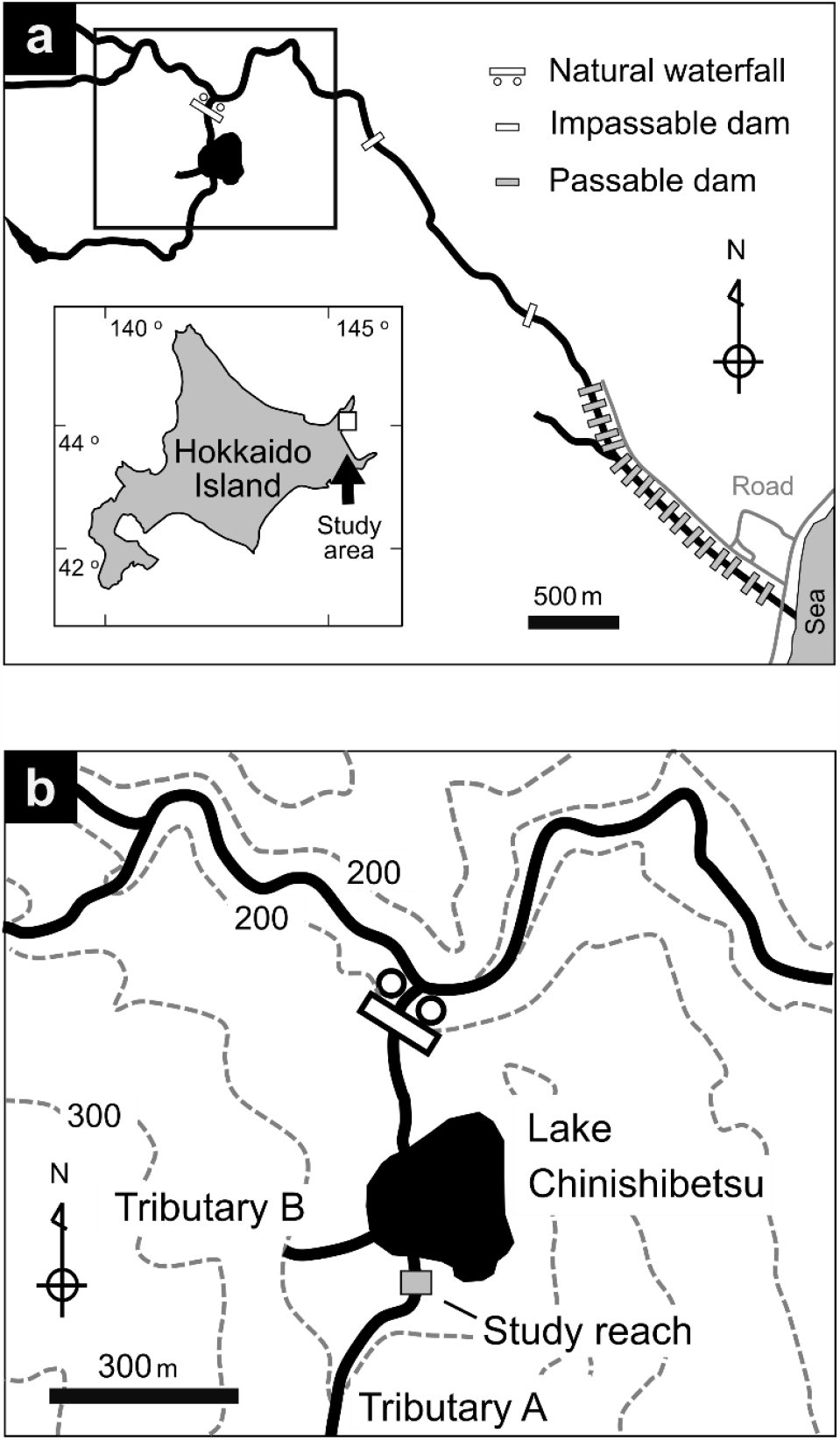
Maps of the study area. (a) Location of the Chinishibetsu River. (b) Locations of the two study inlet tributaries (Tributaries A and B) and Lake Chinishibetsu. Black lines indicate the river’s course, gray lines show roads, and dotted lines show topographic contours. The hatched area indicates the 40-m long study reach established in 2017 to evaluate the population density of rainbow trout in Tributary A.

**Figure 2.**
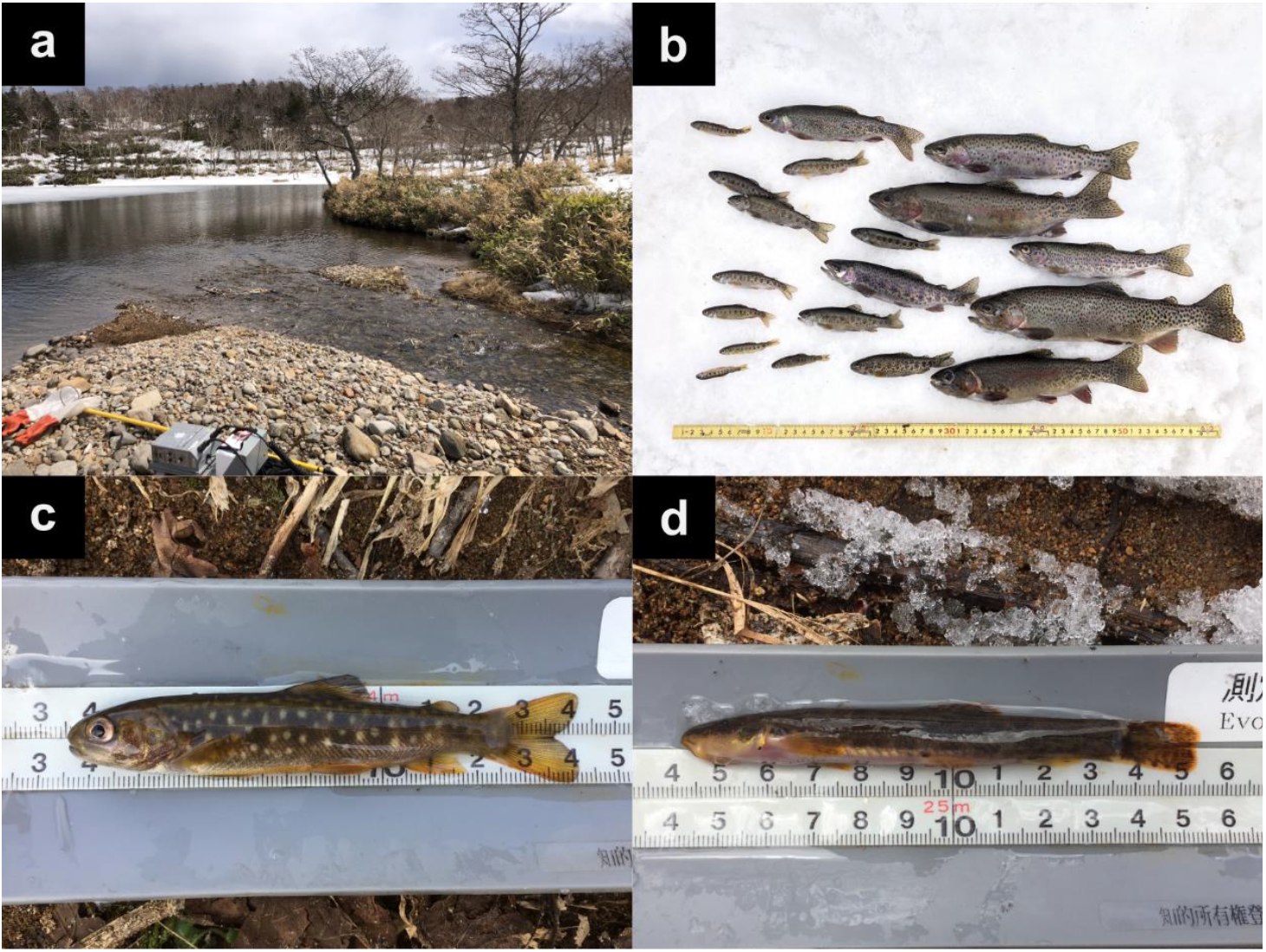
Photographs of the study site and of fish collected in the inlet tributaries of Lake Chinishibetsu. (a) Confluence of Tributary A and Lake Chinishibetsu. (b) Rainbow trout *Oncorhynchus mykiss*. (c) White-spotted charr *Salvelinus leucomaenis*. (d) Siberian stone loach *Barbatula oreas*.

## Results

A total of 104 rainbow trout were collected from the two inlet tributaries of Lake Chinishibetsu (Fig. 2). The sampled trout included 21 immature males, 36 immature females, 42 mature males, and 5 mature females (Fig. 3; Table 1). The density of rainbow trout in Tributary A was estimated at 0.04 ind./m^2^. In addition, 7 white-spotted charr *Salvelinus leucomaenis* and more than 50 Siberian stone loach *Barbatula oreas* were collected in the two tributaries (Fig. 2). However, no southern Asian Dolly Varden charr *Salvelinus curilus*, which are widely distributed in the rivers of the Shiretoko Peninsula, were collected in either tributary.

**Table 1.**
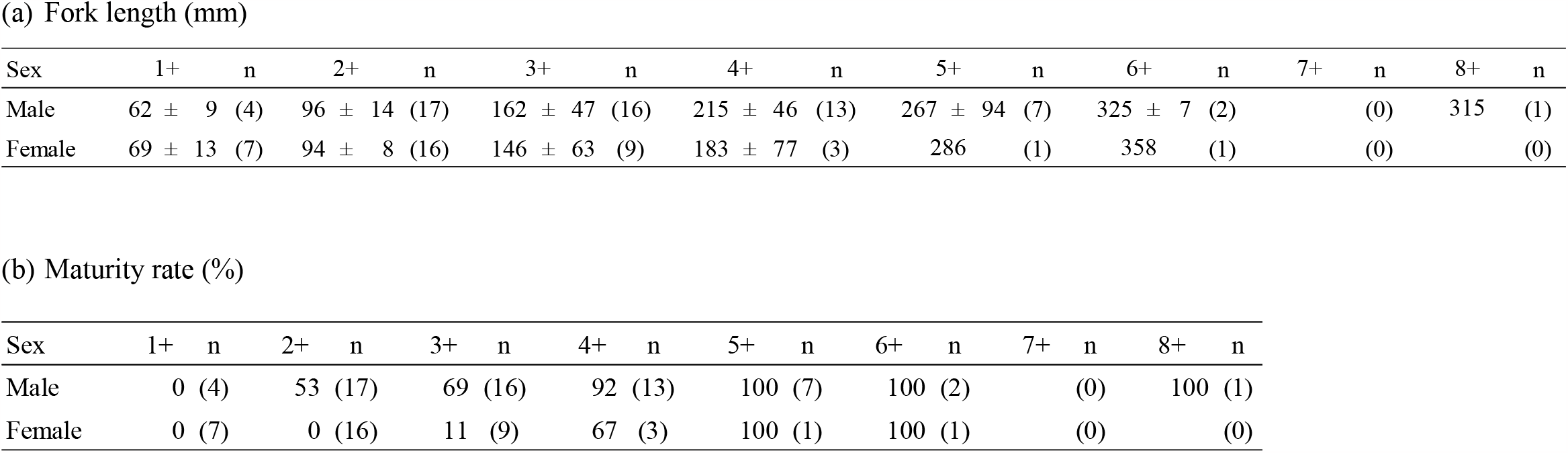
Biotic characteristics of rainbow trout of various ages. Numbers of fish are shown in parentheses.

**Figure 3.**
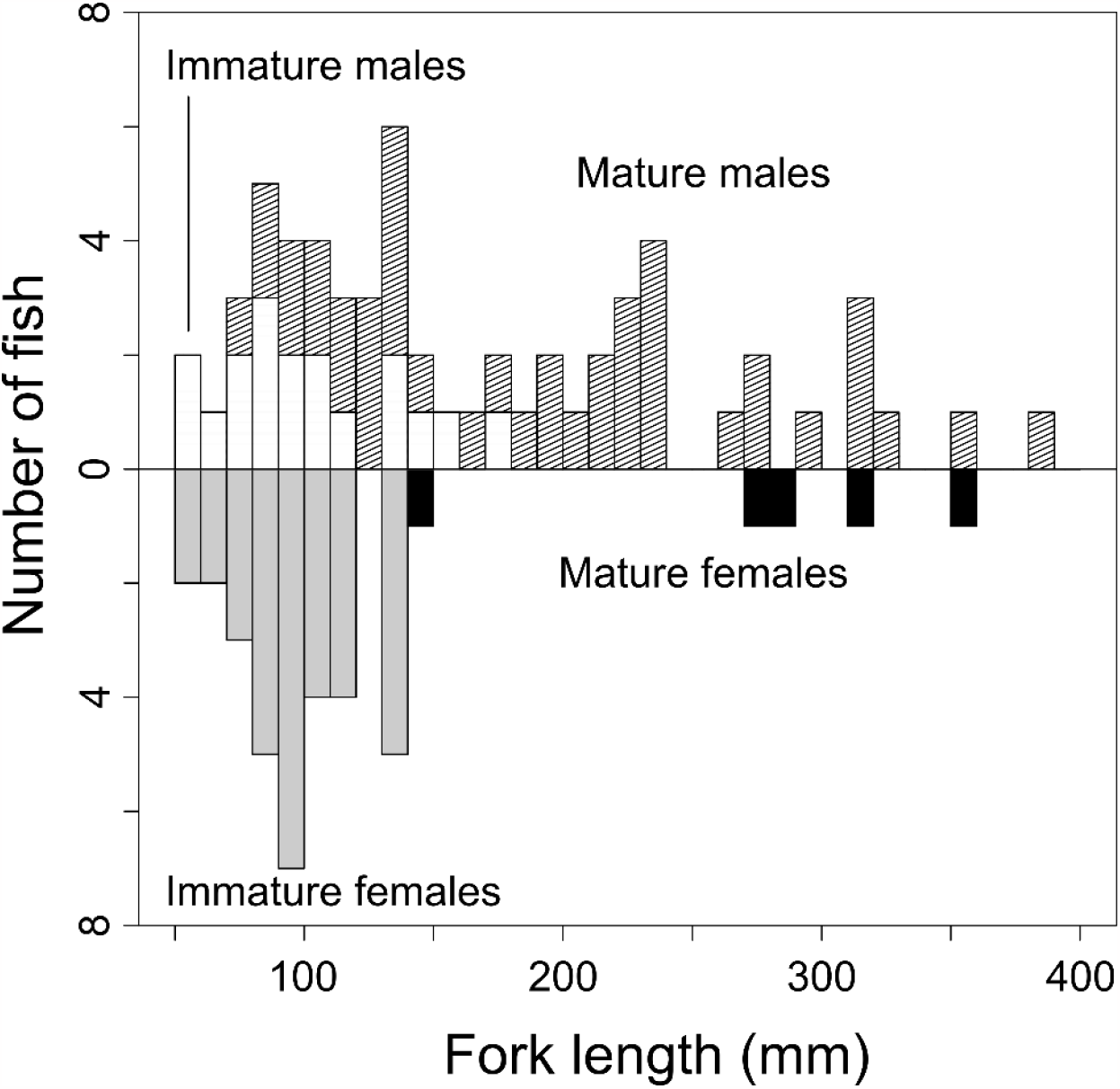
Distribution of body sizes of rainbow trout in the inlet tributaries of Lake Chinishibetsu, Shiretoko Peninsula.

Rainbow trout ranged in FL from 51 to 383 mm (Fig. 3; Table 1). The FL distribution of mature females was bimodal (Fig. 3). Rainbow trout of various ages were collected, ranging from age 1+ to 8+ (Table 1). The oldest rainbow trout was a male aged 8+ (Table 1). Age at first maturity was 2+ for males and 3+ for females (Table 1). The sex ratio of mature rainbow trout was significantly male-biased at 0.89 (Binomial test, *P* < 0.001).

White-spotted charr ranged in FL from 63 to 132 mm (Fig. 4). Ages ranged from 1+ to 3+ years, and all collected fish were immature (Table 2).

**Figure 4.**
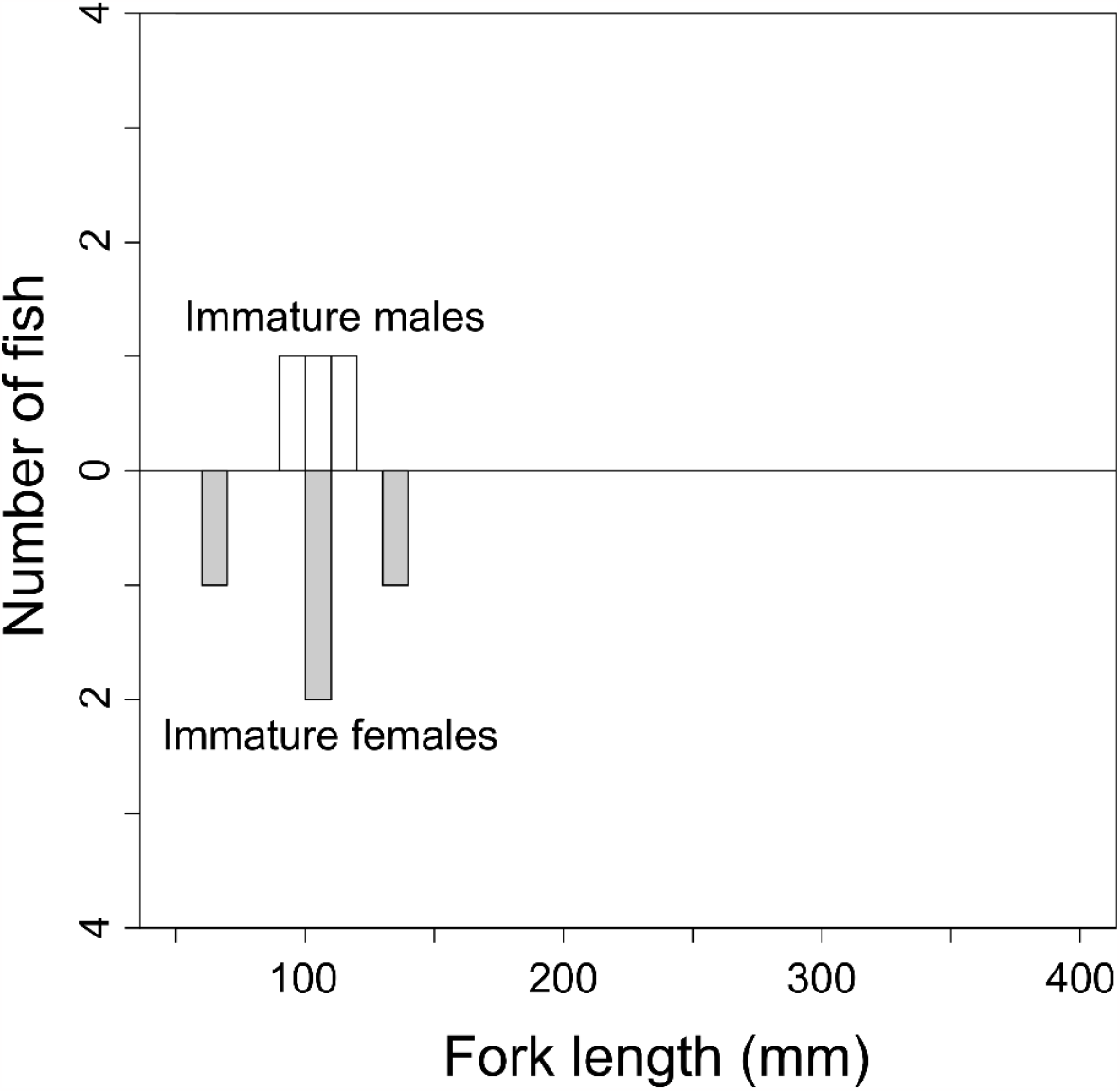
Distribution of body sizes of white-spotted charr in the inlet tributaries of Lake Chinishibetsu, Shiretoko Peninsula.

**Table 2.**
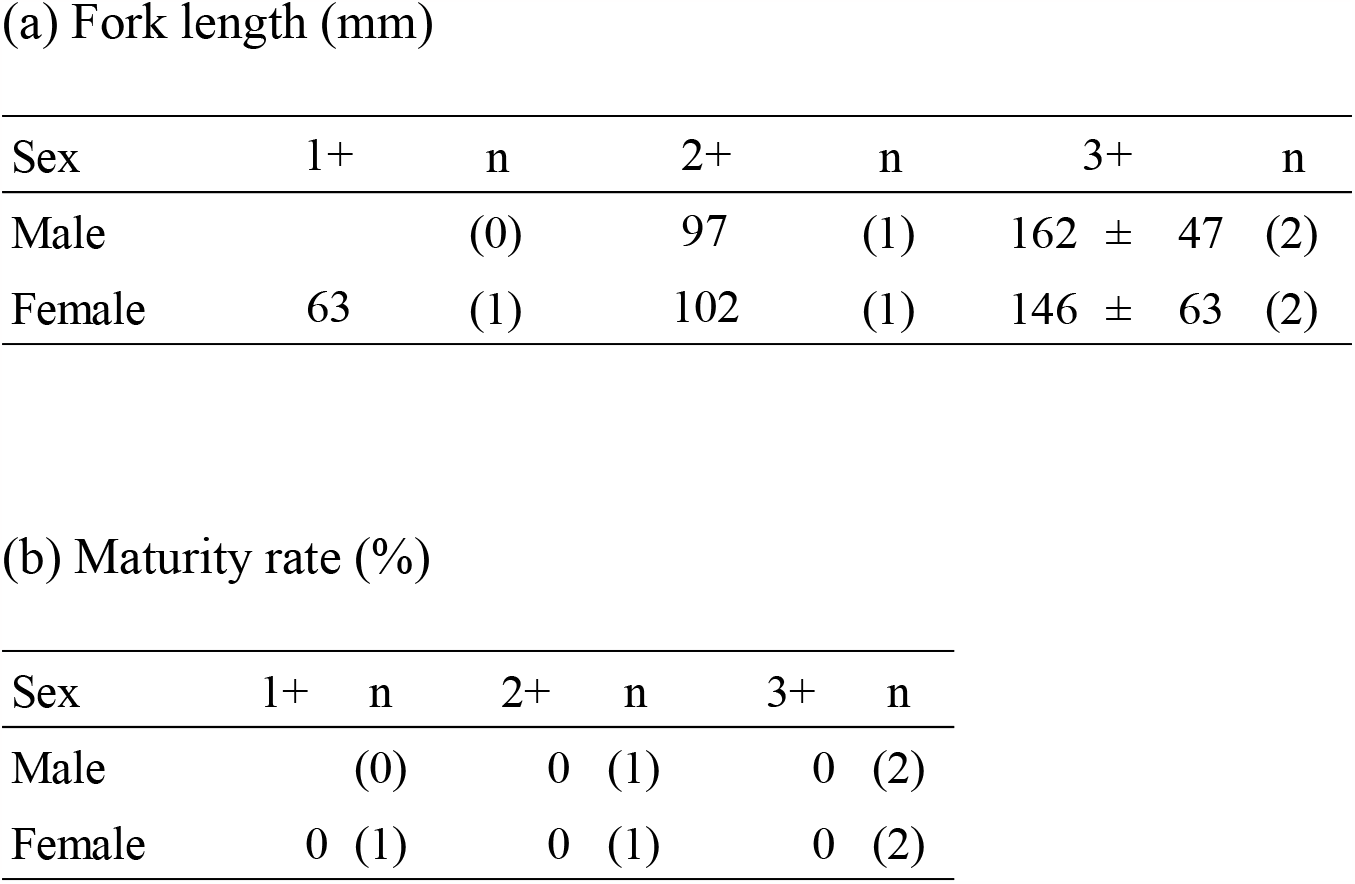
Biotic characteristics of white-spotted charr of various ages. Numbers of fish are shown in parentheses.

## Discussion

Rainbow trout were collected from the inlet tributaries of Lake Chinishibetsu. The collected trout included immature fish, mature males, and mature females and ranged in age from 1+ to 8+ years, including age 1+ individuals that may have been born in the previous year. Additionally, several spawning adults and spawning redds were observed during the study (the author’s personal observation). This suggests that rainbow trout are reproducing in the inlet tributaries of Lake Chinishibetsu.

The sex ratio of mature rainbow trout was significantly male-biased, which is typical of salmonid populations that experience partial migration (Morita et al., 2009; Morita, 2018). In contrast to the middle reaches of the Chinishibetsu River, where migration to lakes and to the sea is blocked by impassable dams and where all rainbow trout collected were smaller than 250 mm FL (Tsuboi et al., 2021), several large individuals (>250 mm FL) were collected in our study. The distribution of body sizes of mature females was bimodal. Taken together, these results indicate the presence of two life-history types, namely river residents and lake migrants, among both sexes in the inlet tributaries of Lake Chinishibetsu.

Siberian stone loach and white-spotted charr were also collected in the inlet tributaries of Lake Chinishibetsu. The presence of natural waterfalls in the outlet river, which prevent fish from migrating upstream, suggests that these fish did not reach the inlet tributaries through natural migration. Artificial expansion of the range of Siberian stone loach can occur as a byproduct of the transplantation of salmonids (Hatakeyama & Kitano, 2018). Additionally, Kishi et al. (2002) have suggested that the Siberian stone loach population in the Chinishibetsu River might originate from past artificial releases. White-spotted charr has been thought to be absent from the Shiretoko Peninsula, including in rivers downstream of the study site (Fausch et al., 1994; Yamamoto et al., 2006; Tsuboi et al., 2021; Hokkaido Regional Forest Office & Forest Realize Co., Ltd., 2022). Therefore, Siberian stone loach and white-spotted charr are likely to have invaded our study area at the same time as rainbow trout.

Only small, juvenile white-spotted charr were collected from the inlet tributaries in our study. In contrast to rainbow trout, which primarily spawns in the spring, white-spotted charr spawns during the fall (Morita et al., 2009; Morita & Morita, 2009; Morita, 2019). White-spotted charr populations that can access lake habitats often include lake migrants, which are individuals born in rivers that subsequently migrate to lakes (Nakano et al., 1990; Yamamoto et al., 1992). In eastern Hokkaido, a large proportion of both male and female charr adopt migratory life histories, and it is uncommon for populations existing upstream of migration barriers to consist solely of river residents (Morita et al., 2009). Therefore, it is likely that larger, mature charr were absent from our sample because they migrated to the lake during the spring when our fish sampling were done in the inlet tributaries.

After the findings of Morita et al. (2003), rainbow trout have subsequently been discovered in the Matsunori River on the Shiretoko Peninsula (Kentaro Morita, personal observation) and in Lake Onne on Kunashiri Island (Kaev & Romasenko, 2021). Kaev & Romasenko (2021) have suggested that the rainbow trout on Kunashiri Island might have strayed from rivers on the main island of Hokkaido. Individuals collected from the Matsunori River and Lake Onne have a silvery body color and easily detachable scales, indicating that they are in good condition for migration to the sea (Kaev & Romasenko, 2021; Kentaro Morita, personal observation). Although expansion of the distribution of rainbow trout through private stocking has been a persistent problem in Hokkaido (Sahashi & Morita, 2016), the possibility of expansion of the distribution via the sea should also be taken into consideration.

The status of several rainbow trout populations on the Shiretoko Peninsula has been monitored through surveys conducted in association with the Shiretoko World Natural Heritage Site. Rainbow trout collected during these surveys, including those conducted downstream of the Chinishibetsu River, are removed. However, despite ongoing removal efforts, rainbow trout continue to be caught downstream of the Chinishibetsu River (Tsuboi et al. 2021; Hokkaido Regional Forest Office & Forest Realize Co., Ltd 2022), and the river remains the primary habitat for rainbow trout on the Shiretoko Peninsula. Therefore, unless rainbow trout are eliminated from Lake Chinishibetsu, which serves as a source of non-native species upstream of the Chinishibetsu River, it will be difficult to control the distribution of this species and to minimize its population on the Shiretoko Peninsula.

## Acknowledgements

We thank Kaori Nakahata for helping with fish measurements in the laboratory, and Kazumasa Ohkuma for obtaining the sampling permit issued by the Governor of Hokkaido. This work was supported by the Global Trout Project funded by the Research Council of Norway [grant no. 287438] and JSPS KAKENHI [grant no. 17K12839]. References include the geographical local names in their titles and bodies.

## References

Crawford, S. S., & Muir, A. M. (2008). Global introductions of salmon and trout in the genus Oncorhynchus: 1870–2007. Reviews in Fish Biology and Fisheries, 18, 313–344.

Dunham, J. B., Pilliod, D. S., & Young, M. K. (2004). Assessing the consequences of nonnative trout in headwater ecosystems in western North America. Fisheries, 29(6), 18–26.

Fausch, K. D. (2007). Introduction, establishment and effects of non-native salmonids: Considering the risk of rainbow trout invasion in the United Kingdom. Journal of Fish Biology, 71, 1–32.

Fausch, K. D., Nakano, S., & Ishigaki, K. (1994). Distribution of two congeneric charrs in streams of Hokkaido Island, Japan: considering multiple factors across scales. Oecologia, 100, 1–12.

Hasegawa, K. (2020). Invasions of rainbow trout and brown trout in Japan: A comparison of invasiveness and impact on native species. Ecology of Freshwater Fish, 29(3), 419–428.

Hatakeyama, R., & Kitano, T. (2018). Reproductive biology of a boreal Nemacheilid loach Barbatula oreas introduced into a temperate river in central Honshu, Japan. Aquaculture Science, 66(2), 123–131.

Hokkaido Regional Forest Office, & Forest Realize Co., Ltd (2022). Report of the 2021 survey on Southern Asian Dolly Varden charr in the Shiretoko Peninsula. p 1–119.

Kaev, A. M., & Romasenko, L. V. (2021). On the capture of rainbow trout Parasalmo mykiss on the Kunashir Island. Journal of Ichthyology, 61(5), 783–786.

Kaizaki, K. (1990). North! A natureling’s travels. Cosmo Books, Tokyo

Kasai F., Yamamoto A., & Mori T. (2010). Age structure and sexual maturity of rainbow trout Oncorhynchus mykiss in Shimatokkari River, Shiretoko Peninsula. Bulletin of the Shiretoko Museum, 31, 7–10.

Kishi, D., Kawaguchi, Y., Kuwahara, T., & Taniguchi Y. (2002). A record of Siberian stone loach Noemacheilus barbatulus toni from Shiretoko Peninsula, Hokkaido. Bulletin of the Shiretoko Museum, 23, 47–50.

Morita, K. (2018). General biology of masu salmon. In: Beamish RJ (ed) The ocean ecology of Pacific salmon and trout. American Fisheries Society, Maryland, p 703–730.

Morita, K. (2019). Trout and charr of Japan. In Kershner JL, Williams JE, Gresswell RE, Lobón-Cerviá, J (eds) Trout and char of the world. American Fisheries Society, Maryland, p 487–515.

Morita, K., Kishi, D., Tsuboi, J., Morita, S. H., & Arai, T. (2003). Rainbow trout and brown trout in Shiretoko Peninsula, Hokkaido, Japan. Bulletin of the Shiretoko Museum, 24, 17–26.

Morita, K., & Morita, S. H., (2007). Alternative life histories and population process of white-spotted charr (salmonid fish). Japanese Journal of Ecology, 57, 13–24.

Morita, K., Morita, S. H., & Yamamoto, S. (2009). Effects of habitat fragmentation by damming on salmonid fishes: lessons from white-spotted charr in Japan. Ecological Research, 24, 711–722.

Nakano, S., Maekawa, K., & Yamamoto, S. (1990). Change of the life cycle of Japanese charr following artificial lake construction by damming. Nippon Suisan Gakkaishi, 56(12), 1901–1905.

Rexstad, E., & Burnham, K. P. (1991). Users Guide for Interactive Program CAPTURE. Colorado Cooperative Fish & Wildlife Research Unit, Colorado State University, Fort Collins, Colorado.

Sahashi, G., & Morita, K. (2016). Potential threat of introduced rainbow trout Oncorhynchus mykiss to native salmonids in the western part of Hokkaido, Japan. Ichthyological Research, 63, 540–544.

Stanković, D., Crivelli, A. J., & Snoj, A. (2015). Rainbow trout in Europe: introduction, naturalization, and impacts. Reviews in Fisheries Science & Aquaculture, 23(1), 39–71.

Tsuboi, J., Morita, K., Sahashi, G., Kuroki, M., Baba, S., & Arlinghaus, R. (2021). Species-specific vulnerability to angling and its size-selectivity in sympatric stream salmonids. Canadian Journal of Fisheries and Aquatic Sciences, 78(10), 1470–1478.

Yamamoto A. (2008). Feeding habits and possible interactions between alien rainbow trout and native salmonids in a small stream on Shiretoko Peninsula, northern Japan. Wildlife Conservation Japan, 11(2), 19–28.

Yamamoto, S., Kitano, S., Maekawa, K., Koizumi, I., & Morita, K. (2006). Introgressive hybridization between Dolly Varden Salvelinus malma and white-spotted charr Salvelinus leucomaenis on Hokkaido Island, Japan. Journal of Fish Biology, 68(A), 68–85.

Yamamoto, S., Nakano, S., & Tokuda, Y. (1992). Variation and divergence of the life-history of Japanese charr Salvelinus leucomaenis in an artificial lake-inlet stream system. Japanese Journal of Ecology, 42, 149–157.

